# Hotspot of *de novo* telomere addition stabilizes linear amplicons in yeast grown in sulfate-limiting conditions

**DOI:** 10.1101/2022.06.12.495855

**Authors:** Remington E. Hoerr, Alex Eng, Celia Payen, Sara C. Di Rienzi, M. K. Raghuraman, Maitreya J. Dunham, Bonita J. Brewer, Katherine L. Friedman

## Abstract

Evolution is driven by the accumulation of competing mutations that influence survival. A broad form of genetic variation is the amplification or deletion of DNA (≥50 bp) referred to as copy number variation. In humans, copy number variation may be inconsequential, contribute to minor phenotypic differences, or cause conditions such as birth defects, neurodevelopmental disorders, and cancers. To identify mechanisms that drive copy number variation, we monitored the experimental evolution of *Saccharomyces cerevisiae* populations grown under sulfate-limiting conditions. Cells with increased copy number of the gene *SUL1*, which encodes a primary sulfate transporter, exhibit a fitness advantage. Previously, we reported interstitial inverted triplications of *SUL1* as the dominant rearrangement in a haploid population. Here, in a diploid population, we find instead that small linear fragments containing *SUL1* form and are sustained over several generations. Many of the linear fragments are stabilized by *de novo* telomere addition within a telomere-like sequence near *SUL1* (within the *SNF5* gene). Using an assay that monitors telomerase action following an induced chromosome break, we show that this region acts as a hotspot of *de novo* telomere addition and that required sequences map to a region of <250 base pairs. Consistent with previous work showing that association of the telomere-binding protein Cdc13 with internal sequences stimulates telomerase recruitment, mutation of a four-nucleotide motif predicted to associate with Cdc13 abolishes *de novo* telomere addition. Our study suggests that internal telomere-like sequences that stimulate *de novo* telomere addition can contribute to adaptation by promoting genomic plasticity.

## Introduction

Genomic variation is the basis of evolution. Mutations, including both single-nucleotide polymorphisms and larger genomic rearrangements, can rise in prevalence within a population if they confer a fitness advantage. Copy number variation (CNV) refers to a genomic rearrangement that results in loss or gain of DNA (Alkan et al. 2011; MacDonald et al. 2014). CNV is most likely to have a phenotypic impact when the DNA that is duplicated or lost contains one or more genes that impact cellular fitness. In humans, CNV is associated with a variety of disorders primarily (but not exclusively) related to intellectual disabilities, developmental delays, and predisposition to cancers (reviewed in Zhang *et al*., 2009; Shaikh, 2017; Doran and Pennington, 2022; R. H. Song *et al*., 2022).

CNV can arise through a variety of chromosomal rearrangements stemming from errors in replication fork progression, recombination, or in DNA damage repair pathways (Hastings et al. 2009). The frequency of CNVs is often increased in cancer cells due to dysregulation of DNA repair or replication pathways (Doran and Pennington 2022). Gene amplification can present in a variety of configurations including both direct and inverted tandem duplication (additional copies at the original location), dispersed duplication (integration of additional copies elsewhere in the genome), or double minute chromosomes that are small circular extrachromosomal fragments of DNA that lack a centromere (Biedler et al. 1983; Chakraborty and Ay 2018; Doran and Pennington 2022).

The origin of CNVs is not well understood, in part because the initiating event may be replaced over time by more beneficial secondary and tertiary events that confer increased growth advantages (Ilić et al. 2022). For example, double minute chromosomes may in some cases serve as recombinational intermediates in the formation of tandem or dispersed duplications (Turner et al. 2017; Rosswog et al. 2021; K. Song et al. 2022). Since most studies of CNV in humans rely on samples from a single point in time, they provide only a snapshot of the most dominant rearrangement (Chakraborty and Ay 2018). This limitation highlights the need for experimental systems in which the temporal relationship between events can be observed.

The budding yeast *Saccharomyces cerevisiae* utilizes DNA repair pathways similar to those of human cells but is more easily manipulated and monitored for genomic rearrangements (Krogh and Symington 2004; Wellinger and Zakian 2012). For example, growth under specific environmental conditions selects for amplification of one of several genes (e.g., *HXT6/7, CUP1, GAP1*, or *SUL1*) (Brown et al. 1998; Gresham et al. 2008; Kao and Sherlock 2008; Gresham et al. 2010; Zhang et al. 2013). Growth of continuous yeast cultures in a chemostat allows the genetic composition of the population to be monitored over time, providing an attractive system to monitor CNV formation and persistence in a heterogenous population. Previously reported evolution experiments with *S. cerevisiae* describe several different types of chromosomal rearrangements that increase gene copy number. Extrachromosomal circular DNAs, similar to the double minute chromosomes described above, have been reported (Gresham et al. 2010). Additionally, persistent linear fragments have also been described (Dorsey et al. 1992; Narayanan et al. 2006). Circular molecules lack DNA ends, but linear fragments must protect their ends from nucleolytic degradation. In most cases, protection of the ends is afforded by the telomere, a repetitive, protein-bound DNA sequence that resists nuclease action and engagement with the DNA repair machinery (Blackburn 1991). Where such linear fragments have been described, they arise from palindromic duplication, resulting in an isochromosomal fragment capped at both ends by copies of the same pre-existing telomere (Dorsey et al. 1992; Narayanan et al. 2006).

In haploid laboratory yeast strains grown in sulfate-limiting chemostats, the fitter variants almost always contain an inverted interstitial triplication of the primary sulfate transporter gene (*SUL1*) and variable amounts of the flanking regions in an otherwise unrearranged chromosome (Araya *et al*. 2010; Payen *et al*. 2014; Brewer *et al*. 2015). Here, we report that diploids of the same laboratory strain appear to follow a different path to *SUL1* amplification. Instead of chromosomally stabilized amplicons, we find extrachromosomal linear fragments that contain *SUL1* and the nearby endogenous telomere at one end and a new telomere added to a sequence centromere proximal to *SUL1* at the other end. Similar fragments arose independently multiple times, suggesting that one or more sequences centromere proximal to *SUL1* have a propensity to stimulate new telomere formation. Indeed, we show that a sequence within the *SNF5* gene is sufficient to stimulate *de novo* telomere addition when moved to an ectopic location. The ability of this sequence to stimulate *de novo* telomere addition requires at least two short sequence motifs that are predicted to associate with the telomere binding protein Cdc13 (Eldridge et al. 2006) and can be functionally replaced by artificial recruitment of Cdc13.

## Methods and Materials

### Strains and Plasmids

Eighteen sulfate-limited chemostat experiments (S701-S712, S10101-3 and S10105-7) were performed on diploid strains of the S288c background (FY3 and FY5) (Gresham et al. 2008; Miller et al. 2013) that were isogenic except for their ARS228 status-*ARS228*/*ARS228, ars228*/*ars228*, and *ARS228*/*ars228*. Creation of the *ars228* mutation has been described (Brewer et al. 2015). All strains for SiRTA analyses were derived from S288c and are listed in Supplementary File 1. Plasmid pAB180 (pRS414-ADHpromoter-GBD-*CDC13*) and pAB220 (pRS414-ADHpromoter-GBD) were gifts of A. Bianchi. Strains for the HO cleavage assay were generated as described in Ngo et al. 2020. In brief, the HO endonuclease recognition sequence was integrated on chromosome VII by one-step gene replacement using plasmid pHO invC (based on JH2017; gift of J. Haber). The *URA3* gene was integrated by one-step gene replacement using pRS306 (Sikorski and Hieter 1989) as template. Putative SiRTAs were inserted on chromosome VII using CRISPR/Cas9 as described (Anand et al. 2017) using gRNA sequence 5’-TGCGGCAAGTTCATCTTCCA. Sequences of different length within *SNF5* were amplified from genomic yeast DNA with appropriate primers (File S1) and additional sequences were added that lie upstream (5’-TTTCTTTGGAAAACGTTGAAAATGAGGTTCTATGATCTAC) and downstream (5’-AGAACATAGAATAAATTTGGTACTGGAACGTTGATTAACT) of the gRNA recognition site. *RAD52* was replaced by one-step gene replacement using pFA6a-TRP1 (Longtine et al 1998) as template.

### Contour-clamped homogeneous electric field (CHEF) gels

Agarose plugs for CHEF gel electrophoresis were generated using the method by Lucas Argueso (described in Kwan et al., 2016). For separation of whole chromosomes, run conditions in the BioRad CHEF-DRII were 1% LE agarose in 0.5XTBE in 2.3 L 0.5% TBE running buffer at 14°C. Switch times were either 47” to 170” at 165V for 39-62 hours or 300” to 900” at 100V for 68 hr. Linear fragments were separated from chromosomes by CHEF gels with switch times 1.7” to 2.6” at 198 V for 20 hr. Restriction analysis of linear fragments was performed by conventional agarose gel electrophoresis in 0.4% ME agarose gels in 1XTBE at 1V/cm for 19 hr at room temperature. Standard protocols for restriction digestion, Southern blotting and hybridization using ^32^P-labeled PCR probes were used. Linear fragments were concentrated from plug supernatants using Centricon Filter devices (Millipore) and then ethanol precipitated.

### Sequence analysis of linear fragments

Whole genome sequencing of fragments purified from plug supernatants and split read analysis were as described (Payen et al. 2014). Briefly, sequencing libraries were generated using Nextera tagmentation (Illumina). Libraries were sequenced on an Illumina MiSeq instrument with paired end 150 bp reads. Number of reads obtained per sample ranged from 42,619 to 345,290 (exact reads for each sample can be found on each data deposit record, see below). Reads were then mapped onto the S. cerevisiae genome with BLAST and filtered for those mapping to chromosome II. Coordinates on chromosome II with large differences in upstream and downstream read coverage were identified as putative junction points. Reads that mapped to these coordinates were aligned to generate junction sequences. Raw sequencing data are deposited at the NIH Sequence Read Archive (SRA) under BioProject ID PRJNA909857.

### HO cleavage assay

The HO cleavage assay was performed as described in Ngo et al. 2020. In brief, cells were grown in SD-Ura medium (+2% raffinose) overnight to an optical density at 600 nm (OD600) of 0.6-0.9. Cells were plated on yeast extract peptone medium with 2% galactose (YEPGal). Simultaneously, cells were serially diluted in water and plated on yeast extract peptone medium with 2% dextrose (YEPD) to determine total cell number. When plasmid selection was required (pRS414, pAB220, and pAB180), cells were plated on synthetic medium lacking tryptophan and containing either 2% glucose or 2% galactose. After 3 days, colonies were counted and at least 100 individual colonies were transferred to plates containing 5-fluoroorotic acid (5-FOA) to determine the frequency of *URA3* loss. For each experiment, at least 30 5-FOA resistant clones were grown in YEPD for DNA extraction by MasterPure™ Yeast DNA Purification Kit (Lucigen). The relative location of each chromosome truncation event was determined using multiplex PCR with primers that anneal centromere or telomere proximal to the SiRTA (File S1). *De novo* telomere addition events were amplified using CHECKCHR7GUIDE1F and TelomereRev as primers. The purified PCR product was sequenced using Sanger sequencing to determine the site of *de novo* telomere addition. Full data sets corresponding to Fig. 3 and 4 are found in Supplementary File 2.

## Results

### Linear fragments containing *SUL1* accumulate in cells grown in sulfate-limiting conditions

Under sulfate-limiting conditions, *S. cerevisiae* cells with increased capacity for sulfate transport exhibit a selective growth advantage (Gresham et al. 2008). The most common genomic rearrangement observed after prolonged competitive growth of haploid lab strains in low-sulfate conditions is an interstitial, inverted triplication of a chromosomal region containing the primary sulfate-transporter gene *SUL1* and its adjacent origin of replication *ARS228*, located ∼25 kb from the right telomere of chromosome II (Payen et al. 2014; Brewer et al. 2015). From the structure of these amplicons (the central copy of the triplication is in inverted orientation) Brewer et al. (2011) proposed a model [Origin Dependent Inverted Repeat Amplification (ODIRA)] involving template switching at preexisting short interrupted inverted repeats that flank the *SUL1* gene and its adjacent origin of replication (*ARS228*) and the *SUL1* gene. This model proposes that, following fork stalling and reversal, inverted repeats at the 3’ end of the leading strands anneal to the complementary repeats on the lagging strand template. Extension of the 3’ ends by DNA polymerase and ligation to the nearest Okazaki fragment creates a closed loop that is subsequently displaced from the replicating chromosome structure (Brewer et al. 2011). Replication of this evicted molecule in the next cell cycle forms a dimeric, inverted circular DNA molecule that can recombine at the *SUL1* locus to generate the interstitial triplication with the inverted central copy (Brewer et al. 2011). The requirement for an origin of replication within the region that is triplicated is supported by the observation that deletion of *ARS228* results in larger amplicons that encompass the next nearest replication origin (Brewer et al. 2015).

A key feature of the ODIRA model is the generation of an extrachromosomal intermediate that later recombines with the chromosomal locus (Brewer et al. 2011). In an attempt to gather evidence for the unique extrachromosomal intermediate, we constructed a diploid heterozygous for the *ARS228* deletion (*ARS228*/*ars228Δ*; Brewer et al. 2015) and two diploid control strains (*ARS228/ARS228* and *ars228Δ/ars228Δ*) and carried out independent evolution experiments in sulfate-limited chemostats (n = 10, 4, and 4, respectively). We reasoned that if the intermediate arose from the *ARS228* chromosome and subsequently recombined with the *ars228Δ* chromosome, the resulting amplicon would contain copies of both *ARS228 and ars228Δ*.

Contrary to expectations, these diploids did not produce the same type of inverted interstitial amplicons of *SUL1* that has been observed in haploid strains. Array comparative genome hybridization (aCGH) of each of the diploid populations after ∼180 generations (33-36 days) under sulfate-limitation revealed consistent amplification of the *SUL1* region (e.g., Fig. 1a, Supplementary Fig. 1). However, in each case the amplicon was not interstitial but continued through the right telomere. The amplified sequences were limited to this region of chromosome II and the new junctions were narrowly clustered in a small region of chromosome II near the *SNF5* gene.

**Fig. 1.**
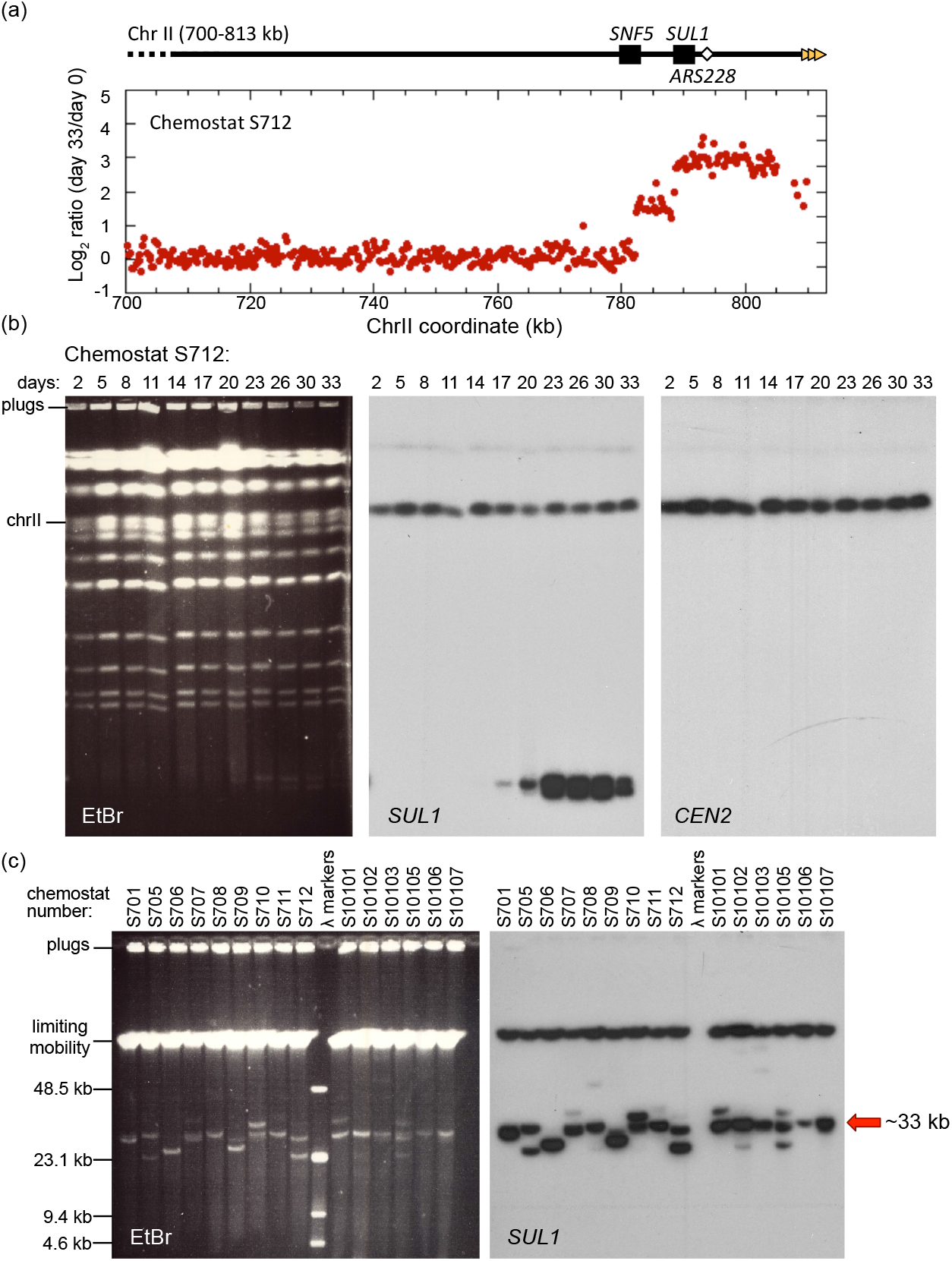
Amplification of *SUL1* chromosomal fragments in sulfate-limited chemostats. a) Array Comparative Genomic Hybridization (aCGH) of the population of a diploid yeast strain (Chemostat run S712) 33 days after continuous growth in a sulfate limited chemostat. The locations of *SNF5, SUL1*, and *ARS228* are indicated in the diagram at the top. The Log2 hybridization ratio of day33/day0 indicates that the right end of chromosome II, containing the gene for the primary sulfur transporter *SUL1*, is present in 8 or more copies in excess of the genomic copies. The presence of two discontinuities at 782 kb and 788 kb suggests two discrete amplicons within the cell population. b) CHEF gel electrophoresis of a single population (S712) sampled across the 33 days of continuous growth in sulfate limited medium. The ethidium bromide-stained gel displays the total yeast karyotype. The CHEF gel was Southern blotted and probed sequentially for *SUL1* and *CEN2* as indicated. One extrachromosomal band of low molecular weight was detected by the *SUL1* probe at day 17 that persists in the population onward. A second, faster migrating band appears at day 23 (more clearly resolved as 2 bands in Figure 1c) and persists in the population onward. c) CHEF gel electrophoresis, under conditions to determine the sizes of extrachromosomal amplicons, from specific days of 15 individual chemostat runs. The ethidium bromide-stained gel and Southern blot hybridized with *SUL1* reveal multiple small extrachromosomal fragments. The sizes of extrachromosomal amplicons vary from 26 kb to 40 kb with a recurring event producing a fragment of approximately 33 kb (red arrow).

To determine the genomic location of the amplicons, we analyzed samples taken every few days through the 33 days of chemostat growth (each day is approximately 6 generations) and analyzed the yeast karyotype by contour-clamped homogeneous electric field (CHEF) gels. Southern blotting and hybridization with probes specific to *SUL1* detected chromosome II as expected, but also revealed the generation of small (∼30 kilobases) molecules that typically were detected at around days 14-17 and often persisted through the end of the chemostat run (Fig. 1b, Supplementary Figs. 1 and 2). Similar sized fragments were present in all samples from all three genotypes (e.g., Fig. 1c, Supplementary Figs. 1 and 2). Because they were generated in diploid cells with both copies of *ARS228* deleted (Supplementary Fig. 1c) we propose that the adjacent origin of replication *ARS229* can support the creation and maintenance of these fragments. The sizes of many of these molecules estimated from their migration on CHEF gels were consistent with the sizes of the amplicon detected by aCGH suggesting that they were acentric linear fragments.

It is notable that the linear fragments often persisted in the population through the end of the experiment. Because the fragments lack a centromere, mitotic partitioning is likely asymmetric. However, the fragments are presumably retained in the population due to the selective growth advantage provided by additional copies of *SUL1*, similar to the maintenance of an ARS plasmid under selection. In half of the cases, chromosomally integrated *SUL1* amplicons were the predominant form by the end of the chemostat run. In many of these cultures, the linear fragments disappeared concurrently with the emergence of new, slower-migrating *SUL1*-hybridizing bands (Supplementary Figs. 1 and 2), suggesting either a precursor-product relationship between the linear fragments and the chromosomally integrated form or the independent occurrence of a second chromosome amplification event conferring increased fitness. One of the most intriguing aspects of these free linear fragments is their consistent small size—ranging between ∼26 and 42 kb (Fig. 1c). Since all of the linear fragments extend through the right telomere of chromosome II, the newly formed left ends of the linear fragments appear to arise near or within *SNF5*, a gene centromere proximal to *SUL1*.

### Linear fragments are stabilized by addition of a new telomere

While selective pressure can partially compensate for the lack of a centromere, persistence of a linear fragment requires protection at both ends from exonucleases. Normal chromosome ends (telomeres) in *S. cerevisiae* consist of a thymine and guanine (TG)-rich strand that extends 10-15 bases past the complementary adenine and cytosine (AC)-rich strand to create a 3’ overhang (Blackburn 1991). This TG-rich single-stranded overhang interacts with the enzyme telomerase, which utilizes its reverse transcriptase catalytic subunit and intrinsic AC-rich RNA template to add telomeric repeats to the chromosome end (Singer and Gottschling 1994; Lingner et al. 1997). In yeast, the 3’ terminating strand consists of a heterogeneous sequence pattern with a single thymine followed by one to three guanines (abbreviated TG1-3) (Wellinger and Zakian 2012).

The aCGH analyses (Fig. 1a, Supplementary Fig. 1) suggested that the right end of the linear fragment contained the endogenous right telomere, but the structure of the left end was initially unclear. More detailed analysis of the linear fragments was facilitated by the development of a method to purify these small molecules through passive diffusion from agarose CHEF gel plugs. Over the course of several weeks, the small fragments diffuse out of the agarose plug and into the supernatant where they can be recovered, while the intact chromosomes remain trapped in the plug (Supplementary Fig. 3). Purified linear fragments were digested with *Eco*NI, which cleaves three times in the terminal region of chromosome II (Fig. 2a). Following separation by CHEF gel, the Southern blotted was probed sequentially with sequences that hybridize to either *SUL1* or telomeric DNA (Fig. 2b). *Eco*NI digestion separated *SUL1* from the right telomere and allowed us to query both ends of the linear fragments. The restriction fragments containing the right telomere hybridized to telomeric DNA (C1-3A/G1-3T) and not to *SUL1* (Fig. 2b, gold rectangles). However, fragments of multiple sizes (ranging from ∼3-25 kb) hybridized to both the *SUL1* and the telomeric probe (Fig. 2b), suggesting that the left end of each fragment was also capped by telomeric DNA.

**Fig. 2.**
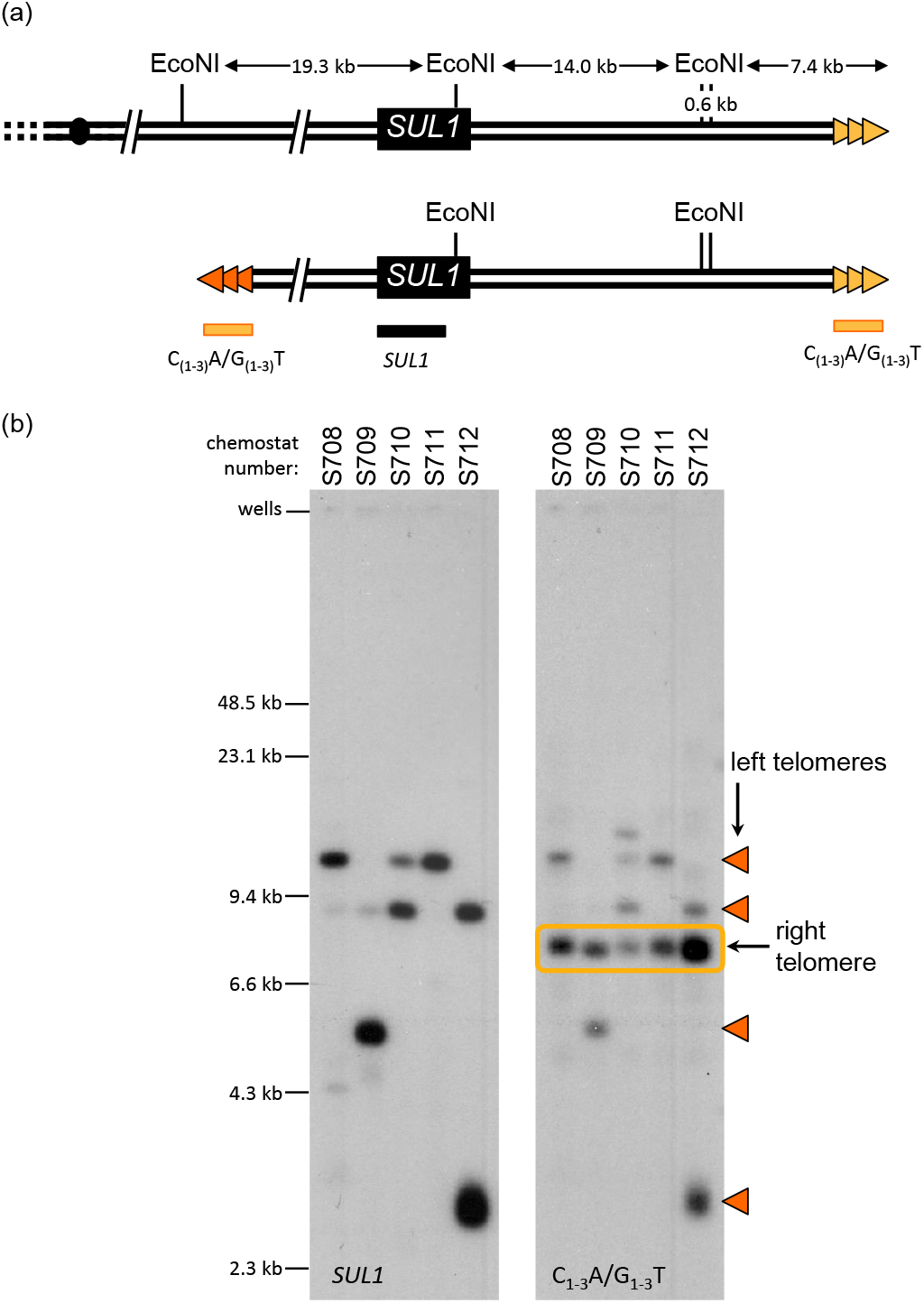
Linear fragments are capped by new telomeric sequences. a) Top: Positions of the *Eco*NI restriction sites at the right end of chromosome II (coordinates 770 to 813 kb). Chromosome II-R telomeric sequences are indicated with gold triangles. Bottom: Structure of linear fragments. Proposed telomeric sequences at variable positions at the left margin of the amplicons are indicated by the orange triangles. Positions of two probes [C(1-3)A/G(1-3)T and *SUL1* (indicated by the horizontal bars)]. b) Linear fragments purified from the plug supernatant (see Supplemental Fig. 3) for five cultures (S708-S712) were digested with *Eco*NI and separated on a CHEF gel. Southern blots were sequentially hybridized with a *SUL1* probe and a telomere specific probe (C(1-3)A/G(1-3)T). Cultures S710 and S712 contain two species of extrachromosomal molecules. Orange triangles indicate the fragments with *de novo* telomeres. Fragments boxed in gold correspond to the native right telomere

**Fig. 3.**
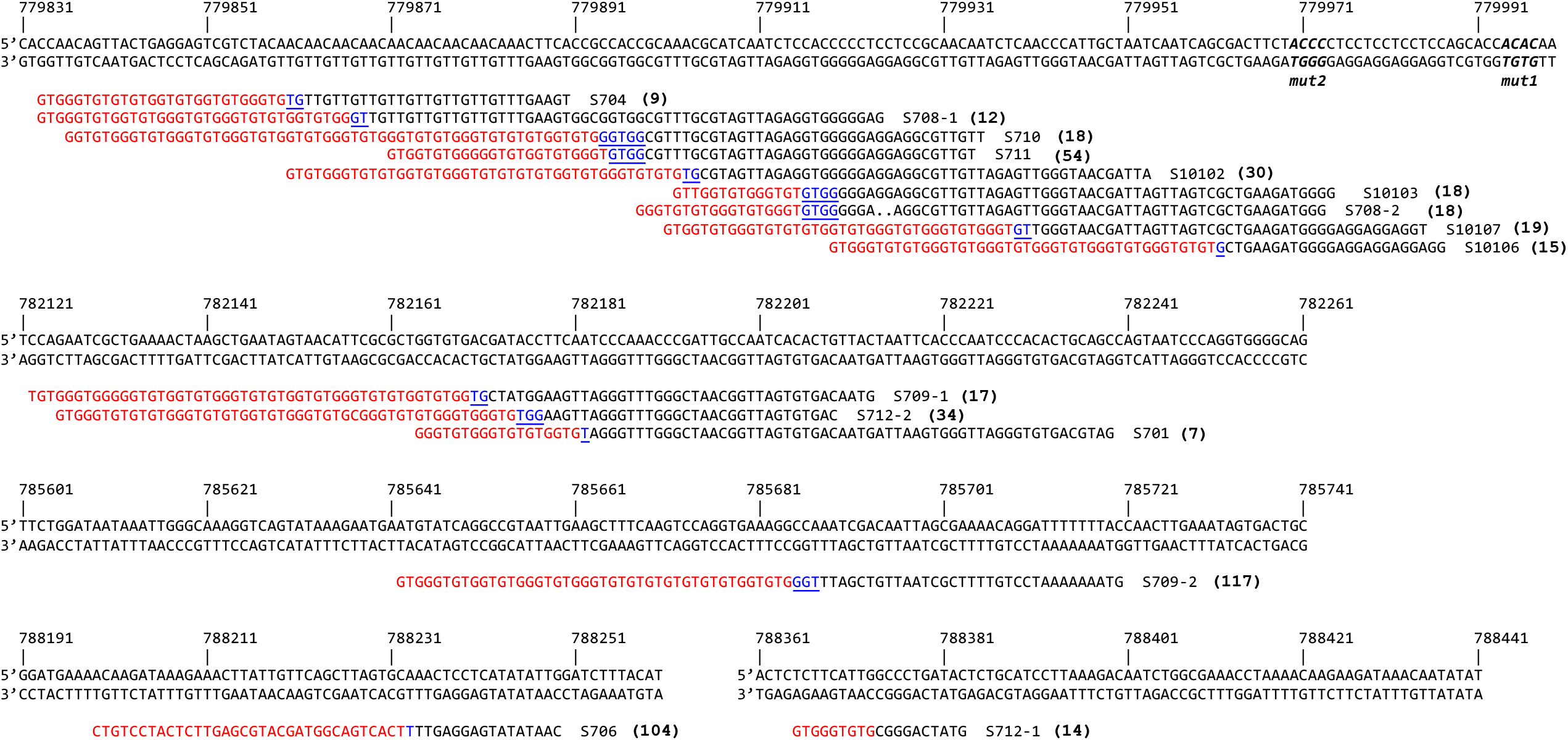
Extrachromosomal fragments purified from plug supernatants were prepared for Illumina sequencing. Split reads (red/black sequences) were aligned to chromosome II (black sequence withchromosome II coordinates). The junctions of the left end of the extrachromosomal fragments each contained a new telomeric sequence (red) and potential seed sequences for telomere addition (blue/underline). The number in parentheses to the right of each sequence corresponds to the number of split reads that mapped to that site. One of the 15 sequences (S706) was a junction with an internal portion of a Y’ element. The remaining 14 contained unique junctions with C(1-3)A/G(1-3)T telomere sequences. The four nucleotide 5’-GxGT-3’ motifs (see Fig. 5) are in bold italics in the (top) reference sequence.

**Fig. 4.**
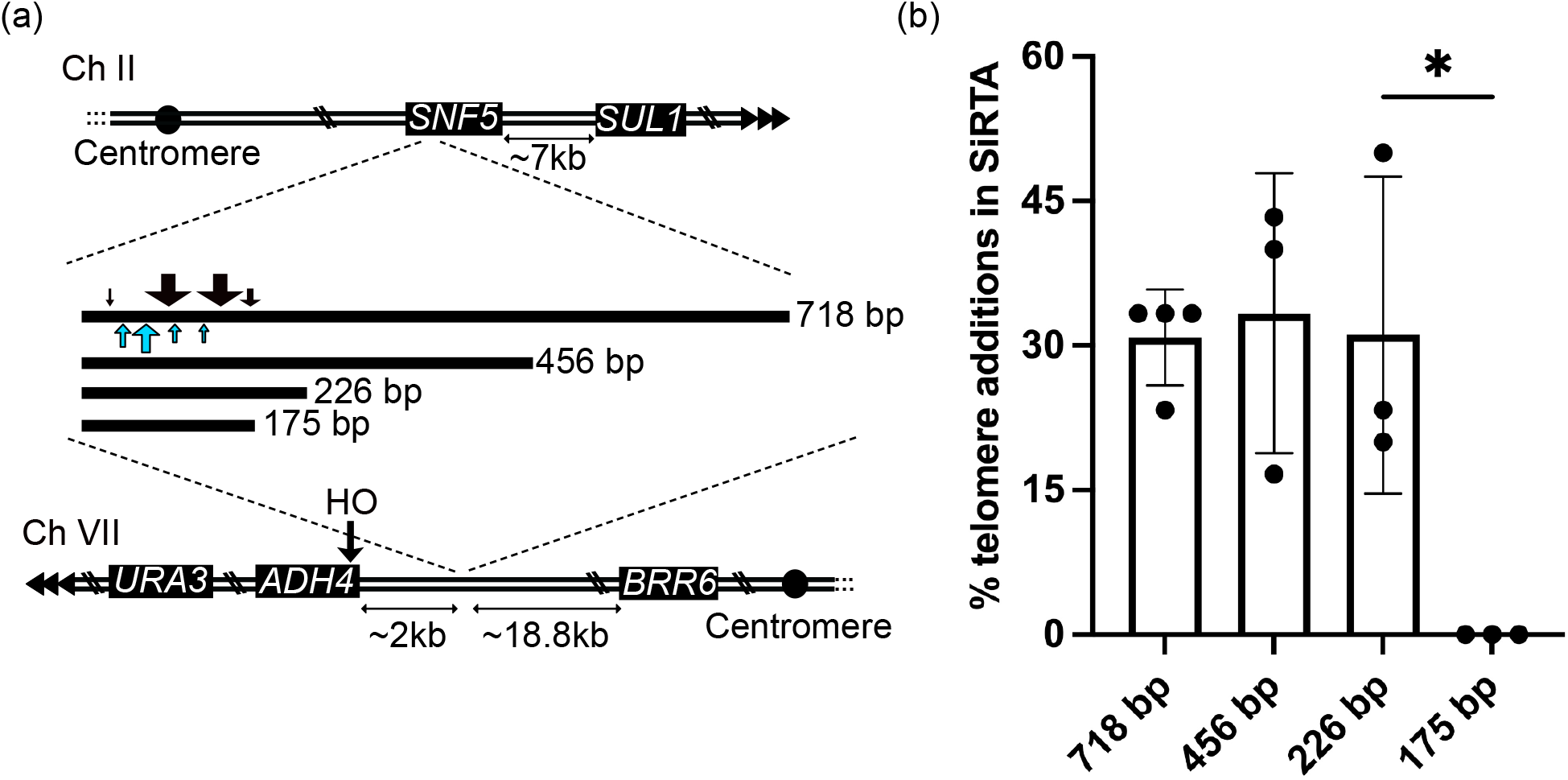
The *SNF5* SiRTA is sufficient to stimulate *de novo* telomere addition at an ectopic location. a) Top schematic: the endogenous right arm of chromosome II; distance between the endogenous *SNF5* sequence and *SUL1* is shown. Triangles represent the endogenous telomere. Middle schematic: lines representing the relative size of each fragment from *SNF5* that was integrated and tested (actual size in bp shown). Black arrows above the 718 bp fragment depict approximate locations of *de novo* telomere addition events stimulated by HO cleavage. Blue arrows below the 718 bp fragment depict approximate locations of spontaneous *de novo* telomere addition events from the chemostat experiments. Arrow size is proportional to the number of independent events at that location (see Supplemental Fig. 5 for exact numbers and locations). Bottom schematic: modified left arm of chromosome VII. Distances between the integrated *SNF5* sequence and the HO cleavage site or the most distal essential gene (*BRR6*) on this chromosome arm are shown. b) Chromosome truncation events following HO cleavage preferentially involve *de novo* telomere addition in the SiRTA. At least 30 clones that survived HO cleavage and lost the *URA3* marker were assayed for each experiment. The percentage of clones that incurred telomere addition within the sequence of interest is shown. Bar height is the average of three to four independent experiments; error bars represent standard deviations. Significant difference is indicated (* p<0.05) by unpaired Student’s *t-test* comparing 226 bp to 175 bp. See File S2 for data used to generate graph.

To identify the sites of telomeric repeat addition, we determined the sequence of fragments purified from the plug supernatants by Illumina sequencing. While short read sequencing is limited in its ability to identify large rearrangements― particularly in organisms with genomes rich in repeated sequences— in yeast, it is perfectly suited for identifying split reads that mark rearrangement junctions. The split reads were confined to the junction between the internal chromosome II sequence and a new telomere— no other junctions were detected along the length of the linear fragment. In addition, the Southern blots and aCGH data were consistent with the absence of large structural changes to the linear fragments. The sequencing results revealed that nine of the 15 fragments contained telomeric repeats clustered within a ∼100 bp region of the gene *SNF5*, approximately ∼7 kb from *SUL1* (Fig. 3). The remaining six events were dispersed across four other regions between *SNF5* and *SUL1* (Fig. 3).

Among the 11 experiments for which we had junction data from both population aCGH and sequence of the *de novo* sites of telomere addition, eight were in perfect agreement. The remaining three identified sequenced junctions that were discordant with the location of the aCGH junction, suggesting an amplicon larger than that predicted by the telomere addition site. These three cases were unique in that the linear fragments had disappeared before the last day of growth—the day on which aCGH was performed. These results are not consistent with the free linear fragments giving rise to the integrated *SUL1* fragments. Rather, they support the idea that a second, independent, more stable chromosomal amplification event (that encompassed a larger section of chromosome II) out competed the linear fragments. Among all of the linear fragments that were sequenced, there was only one example where the new left end had acquired subtelomeric sequences (S706). In all of the other examples the telomeric DNA was appended directly to the chromosome II sequence, consistent with addition of a new telomere to internal sequences by telomerase.

### Sequences in the *SNF5* gene stimulate unusually high levels of *de novo* telomere addition

The observation that many of the telomere addition events associated with extrachromosomal *SUL1* fragments cluster within a small (∼100 bp) region is reminiscent of sequences in the yeast genome that are hotspots of *de novo* telomere addition (termed SiRTAs or Sites of Repair-associated Telomere Addition) (Stellwagen et al. 2003; Obodo et al. 2016; Hoerr et al. 2021). SiRTAs contain strand-specific, TG-rich sequence motifs (Obodo et al. 2016; Ngo et al. 2020). These motifs can provide a substrate for the acquisition of telomeric sequences if they become exposed in single-stranded DNA either as a result of 5’ end resection following a DSB or at nicks in single-stranded gaps. In either case, the resulting TG-rich 3’ overhang can invade a similar sequence and acquire a telomere through break-induced replication or can undergo *de novo* telomere addition by direct action of telomerase (Putnam and Kolodner 2017). Such events are most easily assayed in cases where the new telomere is added to the centromere-proximal side of the break, causing a terminal truncation. In contrast, the putative SiRTA within *SNF5* is oriented in the opposite orientation on the right arm of the chromosome. Rather than stabilizing the centromere-containing region of the chromosome, *de novo* telomere addition stabilizes a terminal acentric fragment containing *SUL1*.

Like telomeres, SiRTAs have an asymmetric sequence pattern in which one strand is relatively rich in T and G nucleotides (Obodo et al. 2016; Ngo et al. 2020). For telomerase to act upon a SiRTA, this TG-rich strand must be exposed in single-stranded DNA. SiRTAs have a bipartite structure consisting of one TG-rich region that is most often the site of telomere addition (the “Core”) and a second TG-rich “Stim” sequence located 5’ to the Core on the TG-rich strand (Obodo et al. 2016). The Stim sequence itself is rarely targeted for telomere addition but is required to stimulate *de novo* telomere addition in the Core (Obodo et al. 2016). Based on this known structure, we speculated that the ∼100 bp region of *SNF5* most frequently observed to undergo telomere addition during generation of the linear fragments represents the Core sequence and that an adjacent sequence would provide Stim function. For a telomere to be added to the left end of the acentric fragment of chromosome II, the TG-rich sequence must be on the bottom or 3’ to 5’ strand (Fig. 3, Fig. 4a). Analysis of this strand showed that a 718 nucleotide (nt) region including and extending 5’ to the putative Core sequence (chromosome II coordinates 779784 to 780501) is unusually TG-rich. Sequences of 300 nt from the 3’ to 5’ strand generated as 10 base sliding windows extending across this region contain, on average, 66% T or G nucleotides [(T+G)/300 = 0.66 #±0.05 n= 42; Supplementary Fig. 4a). In contrast, 300 nt sequences randomly selected from the yeast genome (n=493) contain, on average, 50% T or G nucleotides [(T+G)/300 = 0.50±0.04], as expected from an unbiased distribution relative to each other and the two strands (Supplementary Fig. 4a). To ensure that our analysis included any potential Stim sequence, we initially tested this entire 718 nt sequence for its ability to stimulate *de novo* telomere addition.

To determine the ability of this 718 nt sequence to stimulate telomere addition, we took advantage of an assay in which the sequence of interest is integrated on the left arm of chromosome VII, replacing sequences between chromosome VII coordinates 18086 and 18108 (Ngo et al. 2020). This strain (Lydeard et al. 2010; Obodo et al. 2016) contains a single recognition site for the yeast homothallic switching (HO) endonuclease that allows regulated generation of a double-strand break approximately 2 kb distal to the sequence being tested (Fig. 4a). Of note, the *SNF5* sequence is inserted on chromosome VII in the inverted orientation relative to the centromere compared with its orientation on chromosome II. However, because the endogenous location is on the right arm of chromosome II and the insertion site is on the left arm of chromosome VII, the TG-rich sequence that will stimulate *de novo* telomere addition is, in both cases, located on the bottom (3’ to 5’) strand. In both configurations, exposure of the TG-rich strand in single-stranded DNA (for example, by resection of a DSB occurring to the left of the sequence of interest) followed by telomerase action on that 3’ overhang will stabilize the chromosome fragment located to the right of the lesion. On chromosome II, *de novo* telomere addition will create an acentric terminal fragment. On chromosome VII, *de novo* telomere addition will truncate the chromosome. The site at which the putative SiRTA is integrated is distal to the last essential gene on the chromosome arm (*BRR6*), ensuring that cells adding a *de novo* telomere within the sequence of interest remain viable. Because the sequence being tested is present both at its endogenous location and at the integrated site on chromosome VII, we deleted *RAD52* to prevent repair by ectopic recombination.

Growth of this strain on media containing galactose induces expression of the HO endonuclease, generating a double-strand break (DSB) distal to the putative SiRTA (Lydeard et al. 2010; Obodo et al. 2016). Correct repair restores the cleavage site, leading to a cycle of cleavage and repair that culminates in cell death or mutation/loss of the HO site. Among the cells that survive HO cleavage, those that have lost a *URA3* marker distal to the HO site are identified through selection for growth on media containing 5-fluoroorotic acid (5-FOA) and are subsequently screened through a combination of PCR and Sanger sequencing to identify telomere addition events (Obodo et al. 2016). The fraction of 30 analyzed clones that have undergone *de novo* telomere addition within the sequence of interest represents the “relative” frequency of *de novo* telomere addition and is used as a measure of SiRTA function (Obodo et al. 2016; Epum et al. 2020). Previously described SiRTAs display relative frequencies of *de novo* telomere addition ranging from 10-35% (Obodo et al. 2016; Ngo et al. 2020).

In this case, analysis of 5-FOA resistant clones surviving HO cleavage from four independent assays (30 clones each) showed that 30.8% ±5% had incurred truncation events mapping to the 718 bp sequence (Fig. 4b). Each of the 51 5-FOA resistant clones was analyzed by PCR using one primer that anneals to the AC-rich telomeric strand and a second primer that anneals centromere-proximal to the 718 bp sequence (Mangahas et al. 2001). In every case, a product of the expected size was produced which, when sequenced, revealed the direct addition of telomeric DNA to the chromosome II sequence (File S2). Taken together, these results indicate that this region functions as a SiRTA (subsequently referred to as the “*SNF5*” SiRTA due to its location within the *SNF5* open reading frame). 50 of the 51 telomere addition events mapped to ∼180 bases at the 3’ end of the 718 base region, relative to the TG-rich strand (Fig. 4a, Supplementary Fig. 5). This ∼180 base region includes the ∼100 base region in which telomere addition was observed in the linear fragments and 69% of the HO-induced events map within 3 bases of a telomere-addition event observed on the linear molecules. However, six of nine telomere additions in the linear fragments lie 3’ to the vast majority of telomere additions observed after HO induction (Fig. 3 and Supplementary Fig. 5). It is unclear why events observed in the linear fragments from the sulfate limited chemostats are distributed differently than in the HO assay.

### *De novo* telomere addition requires specific motifs associated with binding of Cdc13

To refine the location of sequences required for *de novo* telomere addition, we tested a series of fragments that included the region within which telomere addition occurred but were progressively truncated at the opposite (5’) end of the TG-rich strand. Fragments of 456 and 226 bp stimulated *de novo* telomere addition as strongly as the full 718 bp sequence (Fig. 4b). In contrast, no telomere addition events were observed in a fragment of 175 bp, despite retention of virtually all sequences that were targeted by telomerase in the context of the larger fragment (Fig. 4a, b). We conclude that the *SNF5* SiRTA has a structure similar to other characterized SiRTAs, with stimulatory sequences that map to the ∼50 bp region delimited by the 226 and 175 bp fragments. We refer to this 50 bp region as the *SNF5* SiRTA Stim. Although the fraction of T and G nucleotides is elevated throughout the 718 bp region (Supplementary Fig. 4b, left axis), the ratio of G to T is notably higher in the centromere proximal 226 nucleotide region that is sufficient for *de novo* telomere addition than in the remainder of the region (Supplementary Fig. 4b, right axis). Because telomeres conform to a pattern of TG1-3, the higher G to T ratio is consistent with a sequence that has the ability to associate with telomeric proteins and/or the RNA template of telomerase.

In *S. cerevisiae*, telomerase is primarily recruited to the telomere by the single-stranded binding protein Cdc13 (Pennock et al. 2001; Bianchi et al. 2004). Binding of Cdc13 to TG-rich Stim sequences rendered single-stranded as a result of 5’ end resection following a DSB has been implicated in the high frequency of *de novo* telomere addition within a SiRTA (Obodo et al. 2016; Epum et al. 2020).Therefore, we hypothesized that the 50 bp *SNF5* SiRTA Stim functions through recruitment of Cdc13. The optimal Cdc13 binding site within telomeric repeats has been characterized (5’-GTGTGGGTGTG) (Anderson et al. 2003; Eldridge et al. 2006), but single nucleotide substitutions within most positions are well tolerated. The largest decrease in Cdc13 binding affinity is observed when G1, G3, or T4 nucleotide positions are substituted (GTGTGGGTGTG), suggesting that a 5’-GxGT-3’ motif may be necessary (but not sufficient) for Cdc13 binding (Anderson et al. 2003; Eldridge et al. 2006). The *SNF5* SiRTA Stim sequence defined above contains two 5’-GxGT-3’ motifs on the TG-rich (bottom) strand (Fig. 5a). To determine if these motifs contribute to the high frequency of *de novo* telomere addition within the SiRTA, we mutated each 5’-GxGT-3’ motif to its complementary CxCA in the context of the 718 bp fragment. Remarkably, mutation of one or both motifs nearly eliminated *de novo* telomere addition (Fig. 5a, b), suggesting that both are required for SiRTA function. This result is consistent with previous reports that SiRTA activity likely requires more than one Cdc13 binding site (Strecker et al. 2017). We hypothesize that the SiRTA sequence must reach a threshold of Cdc13 binding to stimulate *de novo* telomere addition and this threshold is not met in the absence of one or both 5’-GxGT-3’ motifs.

**Fig. 5.**
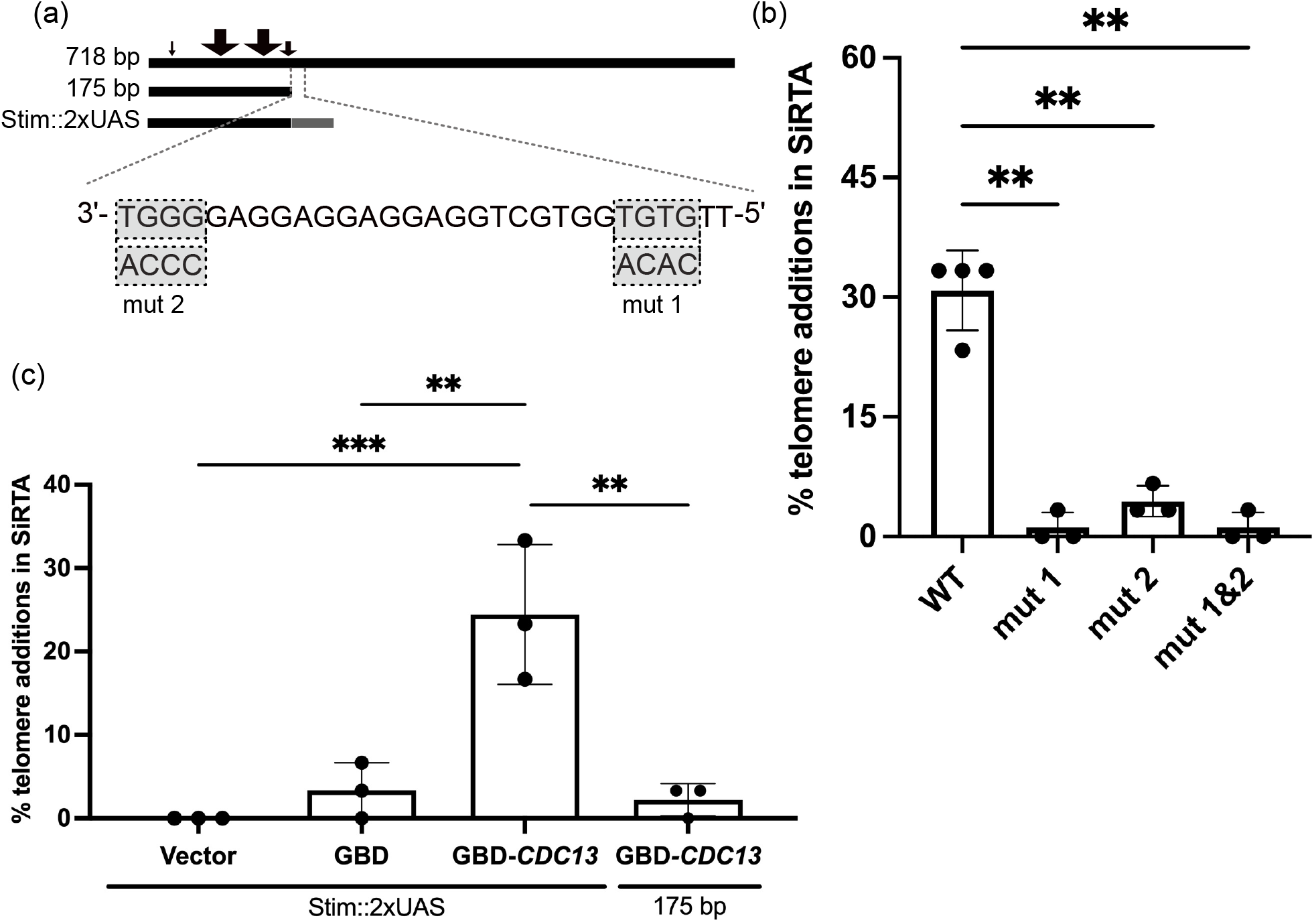
The *SNF5* SiRTA requires Cdc13 binding to stimulate *de novo* telomere addition. a) Top: lines representing 718, 175 bp, and *SNF5* SiRTA Stim::2XUAS fragments integrated and tested on chromosome VII. Sites of *de novo* telomere addition are depicted as in Fig. 4. Bottom: Twenty-nine nucleotides of the TG-rich strand of the *SNF5* SiRTA Stim sequence are shown from 3’ to 5’. Note that a double-strand break that occurs to the left of this sequence will expose the TG-rich sequence in single-stranded DNA following 5’ end resection. 5’-GxGT-3’ motifs and corresponding mutations are highlighted in gray. b) Percentage of 5-FOA resistant clones that incurred *de novo* telomere addition within the 718 bp *SNF5* SiRTA in the presence of the indicated mutation. Wild-type (WT) samples are repeated from Fig. 4 for comparison. Samples with statistically significant values by one-way ANOVA with *post hoc* Dunnett’s are indicated (**, p< 0.01). c) Percentage of 5-FOA resistant clones that incurred *de novo* telomere addition within the 175 bp *SNF5* fragment when 2 copies of the Gal4 UAS are added (Stim::2xUAS) or when the 175 bp fragment is present alone. Cells were transformed with pRS414 (empty vector) or pRS414 expressing either Gal4 DNA binding domain (GBD) or GBD fused to the N-terminus of Cdc13 as indicated. For b and c, at least 30 colonies were assayed for each experiment. Bar height is the average of three to four independent experiments; error bars represent standard deviations. Samples with statistically significant values by one-way ANOVA with *post hoc* Tukey’s are indicated (*, p< 0.01; ***, p<0.001). See Supplementary File 2 for data used to generate graph.

To further interrogate the requirement of Cdc13 binding to stimulate *de novo* telomere addition, we replaced the 50 bp region of the *SNF5* SiRTA Stim sequence with two copies of the Gal4 upstream activating sequence (Stim::2xUAS) to allow specific recruitment of the Gal4 DNA binding domain (GBD) (Bianchi et al. 2004; Zhang and Durocher 2010; Obodo et al. 2016). We introduced an empty vector, a vector expressing GBD alone, or a vector expressing a fusion of GBD to full-length Cdc13 into cells with the *SNF5* SiRTA Stim::2xUAS fragment integrated at the chromosome VII test site. As expected, *SNF5* SiRTA Stim::2xUAS did not support high frequency *de novo* telomere addition with empty vector or GBD alone. Consistent with previous reports that artificial recruitment of Cdc13 to the SiRTA supports *de novo* telomere addition (Bianchi et al. 2004; Zhang and Durocher 2010; Obodo et al. 2016), expression of the GBD-Cdc13 fusion protein supported *de novo* telomere addition in 24.4% #±8.3% of clones tested from three independent assays and stimulation was dependent on the presence of the UAS sequences (Fig. 5c). These results further support our hypothesis that a threshold of Cdc13 binding must be met to stimulate *de novo* telomere addition and that without the 5’-GxGT-3’ motif-containing Stim sequence, this critical threshold is not satisfied.

## Discussion

In this work, we demonstrate that diploid laboratory yeast strains grown in sulfate-limiting conditions generate small, extrachromosomal linear DNA fragments containing the primary sulfate transporter *SUL1* that persist because they provide a selective growth advantage. The linear fragments are capped at one end by sequences from the endogenous right telomere and at the other end by *de novo* telomere addition events clustered in a region centromere proximal to *SUL1*. Fragments of similar, but not identical, size arise in multiple independent cultures, suggesting that some sequence feature or features in this region are responsible for the creation and/or stabilization of these linear fragments. These results raise three intertwined questions: 1) What features of the region make it a hotspot for telomere addition? 2) What process generates the substrate for telomere addition in the sulfate limiting chemostats (in the absence of HO cleavage)? And 3) Why do the diploids produce a different form of *SUL1* amplicon than their isogenic haploids?

### Telomere addition

We find multiple sequence features within the *SNF5* gene that contribute to the stabilization of linear fragments through *de novo* telomere addition. We demonstrate that a DNA fragment from the *SNF5* gene that contains the common sites of telomere addition is sufficient to stimulate *de novo* telomere addition at an ectopic location, similar to other characterized hotspots of *de novo* telomere addition (SiRTAs) (Obodo et al. 2016; Ngo et al. 2020). Previous work demonstrated that association of Cdc13 with a SiRTA is critical for the stimulation of telomere addition (Obodo et al. 2016; Epum et al. 2020). However, the number and/or affinity of Cdc13 binding sites sufficient to support *de novo* telomere formation are unclear. Strecker *et al*. (2017) placed differing lengths of telomeric repeats adjacent to a double-strand break to determine a threshold length above which telomere addition is stimulated. Telomere addition is strongly inhibited at sequences below that threshold length (Strecker et al. 2017). Hypomorphic mutations in Cdc13 increase this threshold value, suggesting that the net number and/or affinity of Cdc13 binding sites likely determines the threshold at which a sequence acts to stimulate telomere addition (Strecker et al. 2017). Our results are consistent with this hypothesis.

Remarkably, mutation of just four nucleotides corresponding to a predicted binding site for Cdc13 (Anderson et al. 2003; Eldridge et al. 2006) nearly eliminates detectable *de novo* telomere addition at the *SNF5* SiRTA in the ectopic assay. Our results suggest that there are likely two Cdc13 binding sites in the ∼50 bp fragment that together constitute the *SNF5* SiRTA Stim. Mutation of either site lowers the total amount of Cdc13 binding below the threshold required for this sequence to function as a hotspot of *de novo* telomere addition. Consistent with this interpretation, replacement of the 50 bp *SNF5* SiRTA Stim sequence with two copies of the Gal4 upstream activating sequence supports *de novo* telomere addition only when cells express the Gal4 DNA binding domain fused to Cdc13.

### Substrate for telomere addition

We showed that *SNF5* SiRTA activity is supported by a minimal fragment of 226 bp. This small region comprises approximately ∼1% (226 bp/20.2 kb) of the total region of chromosome VII in which a viable chromosome truncation could occur (between the HO site and the most distal essential gene on this chromosome arm), and yet nearly one third of all truncation events were mapped to the *SNF5* SiRTA. This clustering of *de novo* telomere additions in a small region of *SNF5* could result from chromosome fragility in this region; however, observations from the ectopic assay argue against this model. The *SNF5* SiRTA undergoes unusually high levels of *de novo* telomere addition even when the initiating chromosome break is intentionally induced two kilobases distal to the eventual site at which telomerase acts, suggesting that it is a site at which DNA ends are preferentially repaired. We conclude that this region of *SNF5* is not necessarily more prone to double stranded breaks, but rather has the fortuitous combination of features that permit telomere addition.

While we have not detected double stranded breaks in chemostat cultures, we have evidence that replication errors, instead of double stranded breaks, may generate aberrant intermediates that are subsequently targeted for repair (Brewer *et al*. 2015). The discontinuous nature of DNA replication forks provides an opportunity for nascent leading strands to anneal to the lagging strand template during fork reversal at sites of interrupted inverted repeats (described as ODIRA in Brewer et al. 2011). In a scan of the region at the 5’ end of the *SNF5* gene we find a ∼5 fold increase in the density of short interrupted inverted repeats (Supplemental Fig. 6a) resulting primarily from a sequence in the promoter region (5’-TGTTGTTG-3’) that is complementary to the several stretches of CAA triplet repeats within the N-terminal coding region of *SNF5* (Fig. 6a). When compared to >10,000 regions across the genome, the *SNF5* region is enriched for short interrupted inverted repeats (Supplemental Fig. 6b). The regression of a replication fork proceeding toward the centromere from *ARS228* or another distal origin (e.g., *ARS229*) could provide multiple opportunities for this sequence on the leading strand to find a match with a complementary sequence in the single stranded gaps on the lagging strand and thereby create a “closed” fork (Fig. 6b). After completion of replication by a fork established centromere proximal to the *SNF5* region, the hairpin capped linear could be released (Fig. 6b) (Brewer et al. 2011). Processing of the single-stranded loop by nucleases provides the appropriate substrate for telomere addition (Fig. 6c, d).

**Figure 6.**
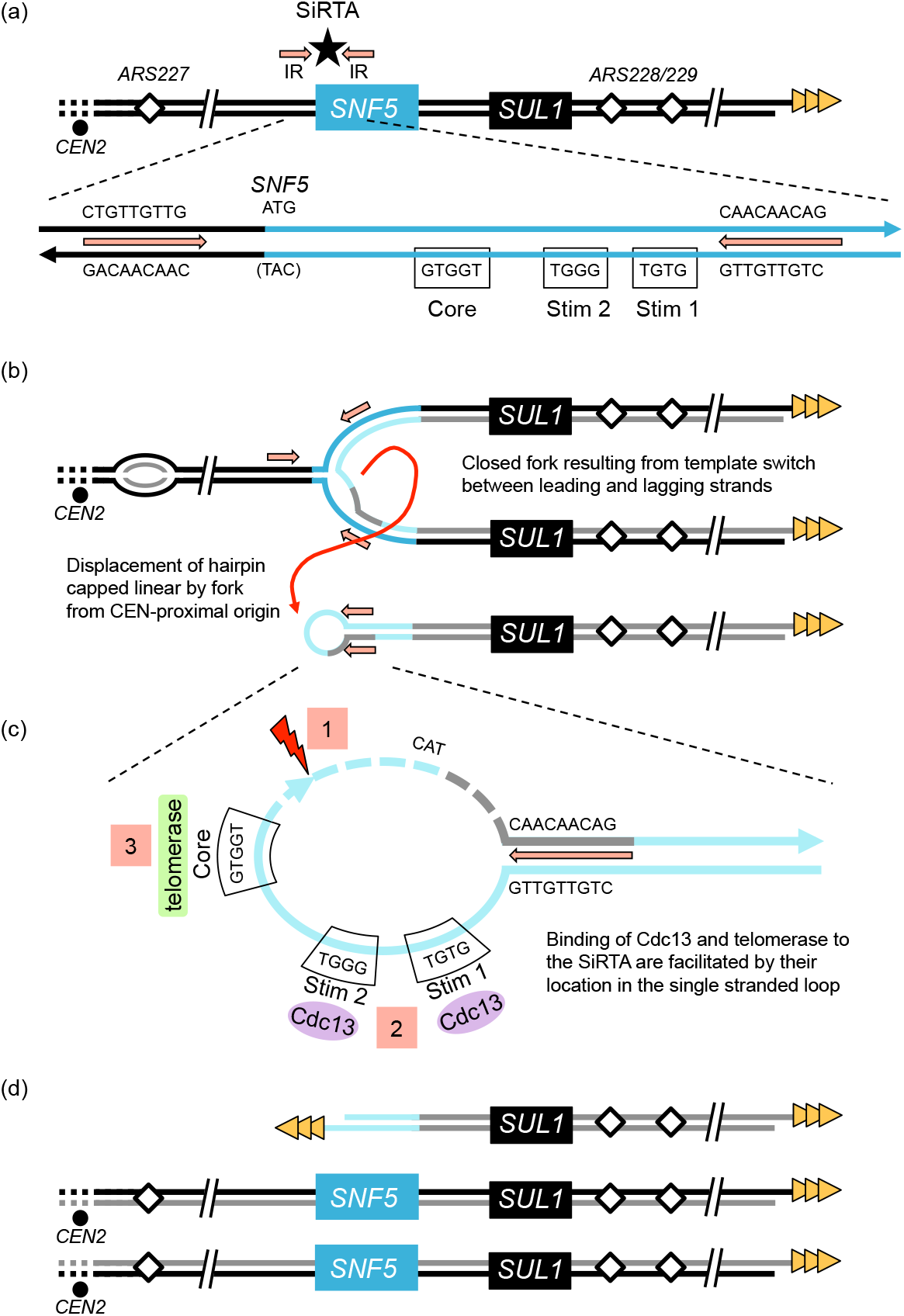
A model to explain how an ODIRA event may generate a substrate for *de novo* telomere addition at the *SNF5* SiRTA. a) The endogenous right arm of chromosome II contains interrupted inverted repeats (IR; orange arrows) flanking the *SNF5* SiRTA (star) and the TG-rich strand of the right telomere (gold triangles). In the expanded schematic the relevant sequences are shown for one possible pair of inverted repeats (orange arrows), the start codon of *SNF5* (ATG), the Core of the *SNF5* SiRTA, and the two 5’-GxGT-3’ motifs from Figure 5 (labeled Stim 1 2). b) Ligation of the leading strand to the lagging strand following fork reversal at the leftward moving fork and template switching at the inverted repeats results in a “closed” fork with the rightward moving fork completing replication through the right chromosomal telomere. When a replication fork from *ARS227*, proceeding towards the telomere, encounters the closed fork structure, a hairpin capped linear is displaced (red curved arrow). c) In the displaced hairpin-capped linear the inverted repeats form the base of the terminal single stranded loop that contains the SiRTA Stim and Core sequences. Cleavage of the hairpin to the 3’ side of the SiRTA (1) and resection at the site of the break (dashed lines) exposes a 3’ single stranded tail containing SiRTA sequences which can be bound by Cdc13 (2) and telomerase (3) to form a *de novo* telomere. d) In this model, after a single round of replication, the chromosome that experienced the closed fork produces two unaltered sister chromatids and the linear fragment containing *SUL1* which is now protected by a *de novo* telomere addition at the *SNF5* SiRTA. (Normal replication of the homolog in this diploid is not shown.) Under sulfate-limiting conditions, the cells containing the linear fragment are retained in the population due to the growth advantage provided by one or more extra copies of *SUL1*.

### Ploidy specific amplification pathways

In previous experiments with haploid laboratory strains, we have not observed the generation of linear fragments—essentially all events were interstitial inverted triplications on chromosome II (Payen et al. 2014). In some cases, rearrangements that would be lethal in haploids are tolerated in diploids by the presence of a second, unrearranged chromosome (Zhang et al. 2013). It is unlikely that a similar phenomenon is affecting the nature of *SUL1* amplification in diploid and haploid cells as there are no essential genes beyond *SNF5* on chromosome II, so loss of this terminal fragment should be tolerable in haploids. In point of fact, we do not detect loss of the distal markers on chromosome II in the diploids— both homologs of chromosome II remain intact. The aCGH profiles indicated that there were no regions at the right end of chromosome II that were present at fewer than two copies per diploid genome and CHEF gels did not reveal any truncated variants of chromosome II (Fig. 1 and Supplemental Fig. 1, and 2). While these results are consistent with repair of a broken chromosome II by homology directed repair off the homolog or sister chromatid (perhaps by break-induced replication), it is also possible that a double stranded break was not the initiating event. We would like to propose that the same type of ODIRA event that can explain the events that occur in haploid cells is also occurring in the diploids, but that the resolution of the intermediate differs between haploids and diploids. Differences in the regulation of repair pathways between haploid and diploid cells in this laboratory strain may contribute to the different outcomes (Li and Tye 2011). There may be a race between replication of the hairpin intermediate and the processing of the single stranded loop that is influenced by the levels of repair enzymes. In support of this notion, we have recently observed that in some haploid wild strains with varied genetic backgrounds, linear amplicons similar to those described here, also arising from the vicinity of the *SNF5* gene, are the preferred amplicon (manuscript in preparation). Analysis of the transcriptomes and proteomes of these strains could identify co-varying RNA/proteins that influence the amplicon outcome. Regardless of mechanism, our observations in yeast may provide a useful model for the types of rearrangements that occur in humans and other organisms. Finally, results presented here suggest that interstitial TG-rich hotspots of *de novo* telomere addition that are oppositely oriented relative to the chromosome arm can stabilize an acentric fragment to provide, in some circumstances, a selective advantage. These observations highlight a possible evolutionary consequence of SiRTA sequences regardless of orientation, by increasing adaptation and genome plasticity.

## Supporting information

Supplementary figures Hoerr et al.

## Data Availability

Strains and plasmids are available upon request. Source data for Figs. 4, 5, and Supplemental Fig. 5 are provided in Supplemental File 2. Sequencing data are available from the NIH Sequence Read Archive (SRA) under BioProject ID PRJNA909857. aCGH data are available from the NIH Gene Expression Omnibus (GEO) under accession GSE220549.

## Acknowledgements

We thank James Haber and Alessandro Bianchi for generously providing us with strains and plasmids and Natalie Guzman for initial exploratory studies.

