## Supplementary figures Hoerr et al. for "Hotspot of *de novo* telomere addition stabilizes linear amplicons in yeast grown in sulfate-limiting conditions"

Figure S1 (a)

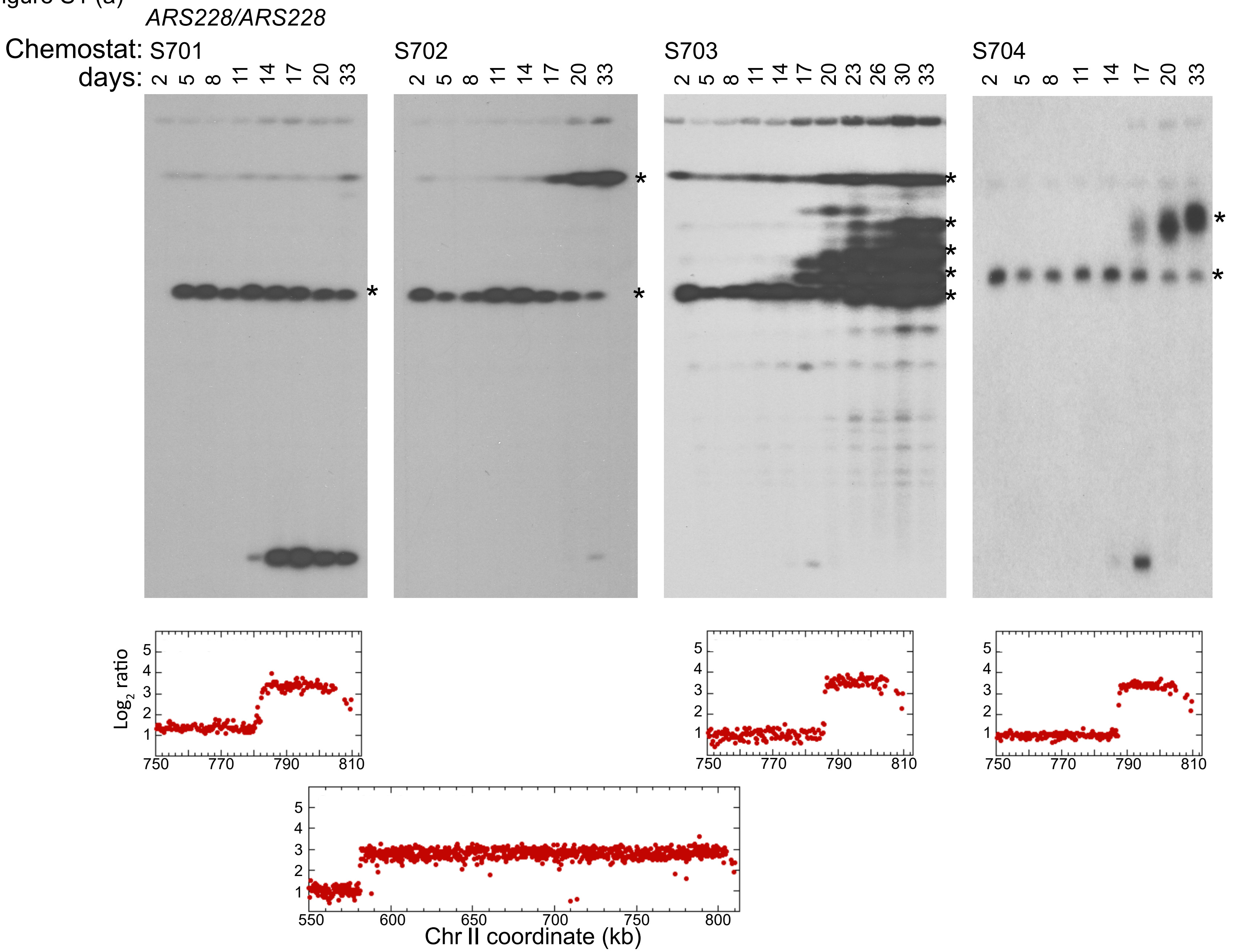

ARS228/ARS228
Chemostat: S701
days: 2 5 8 11 14 17 20 33
S702
2 5 8 11 14 17 20 33
S703
2 5 8 11 14 17 20 23 26 30 33
S704
2 5 8 11 14 17 20 33
Log2 ratio
Chr II coordinate (kb)

Figure S1 (b) *ARS228/ars228*

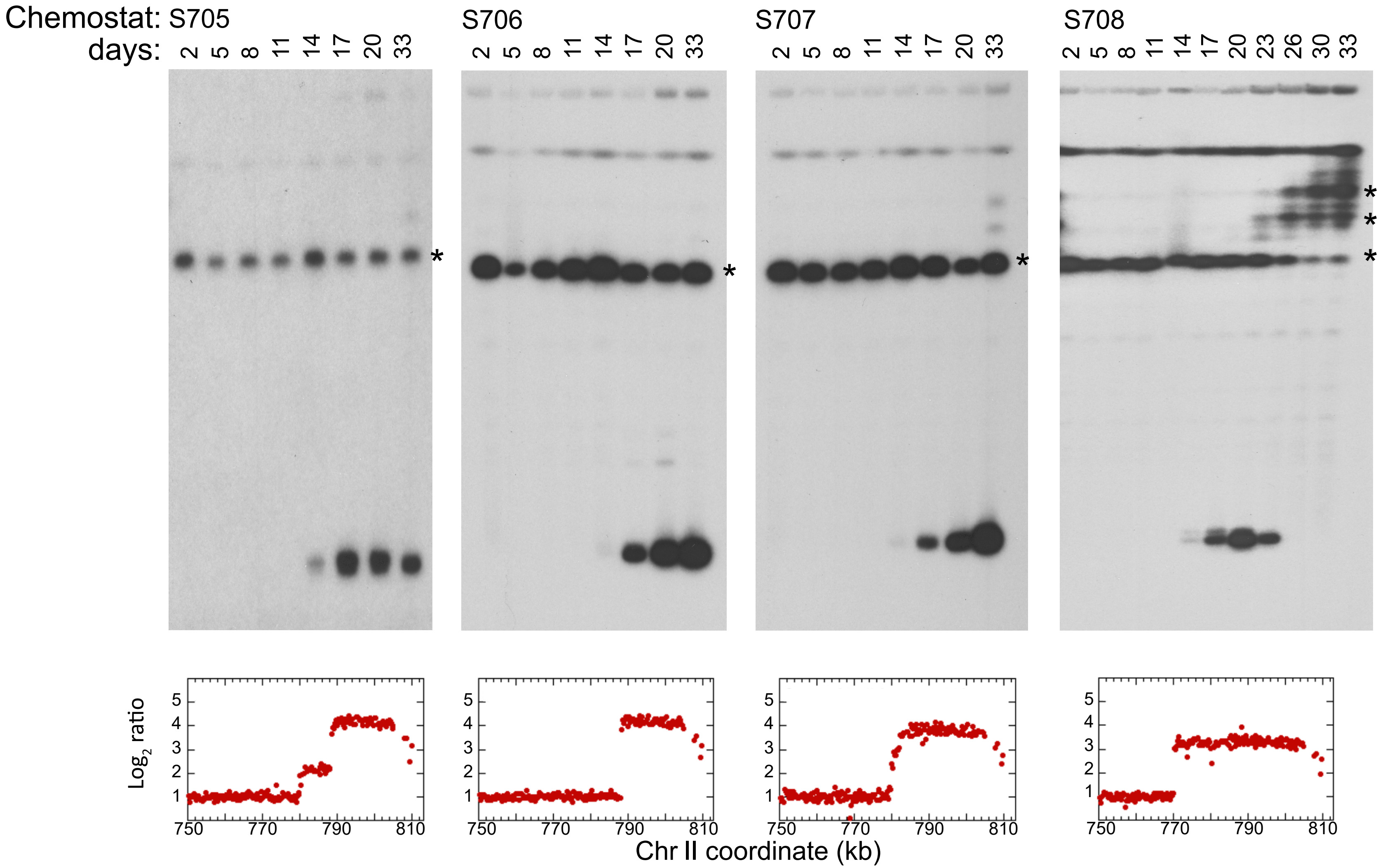

Figure S1 (c)

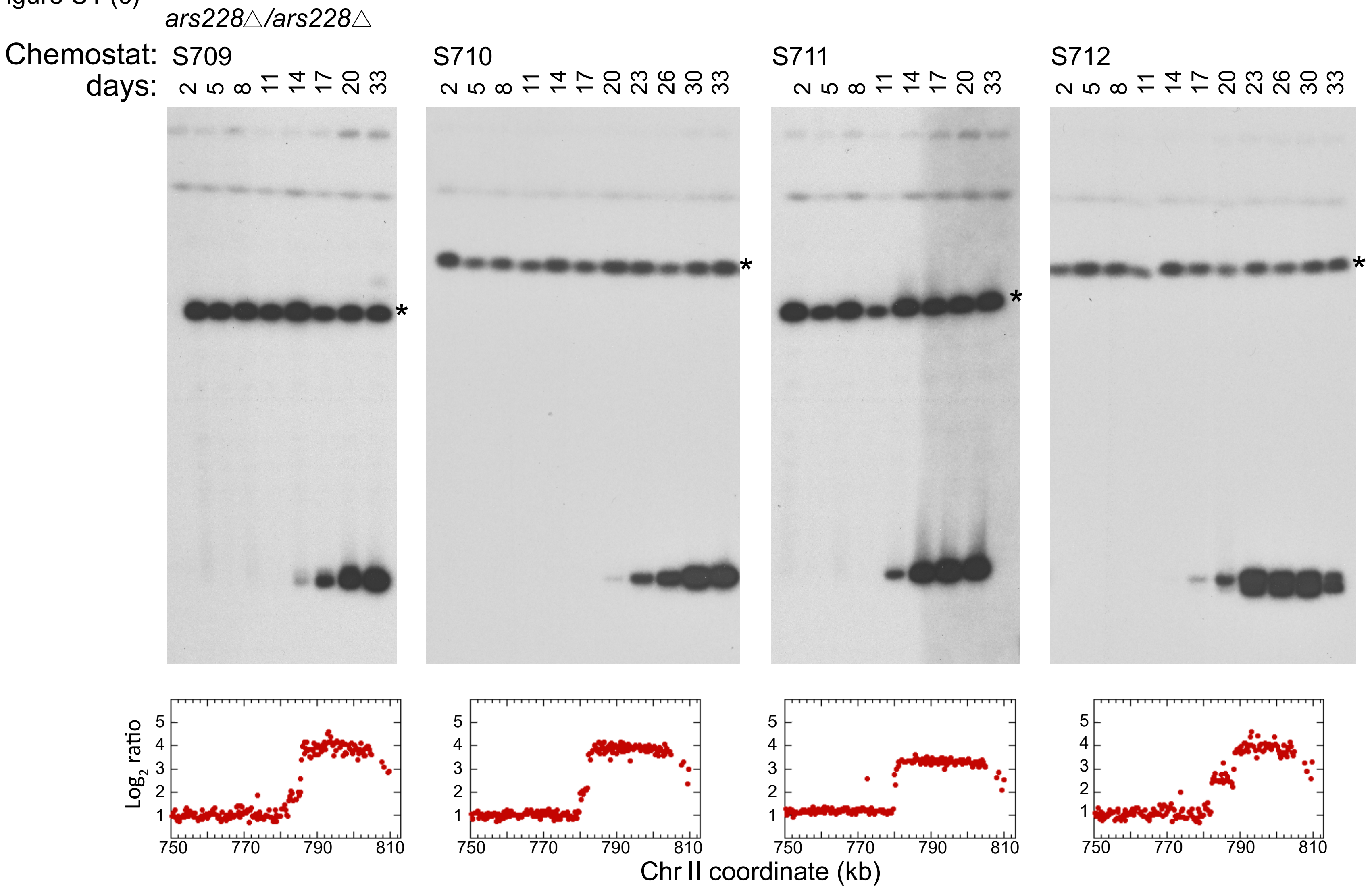

*ars228△/ars228△*
Chemostat: S709
S710
S711
S712
days:
2 5 8 11 14 17 20 33
2 5 8 11 14 17 20 23 26 30 33
2 5 8 11 14 17 20 33
2 5 8 11 14 17 20 23 26 30 33
*
Log2 ratio
1
2
3
4
5
750
770
790
810
Chr II coordinate (kb)

Figure S1. CHEF gel and aCGH analysis of isogenic diploid strains differing only at the *ARS228* locus. (a) A*RS228/ARS228* diploids; (b) *ARS228/ars228Δ* diploids; (c) *ars228Δ/ ars228Δ* diploids. Specific days of the continuous culture were examined based on preliminary sampling to determine the initial appearance and disappearance of the extrachromosomal amplicons. Hybridization with the *SUL1* probe is shown. Bands hybridizing with a *CEN2* probe upon stripping and reprobing of the blots are indicated with an (*). Population samples were harvested from day 33 and processed for aCGH. Amplified regions were limited to the right end of chromosome II. The terminal 65 kb of chromosome II are shown for all but one culture (S702).

Figure S2

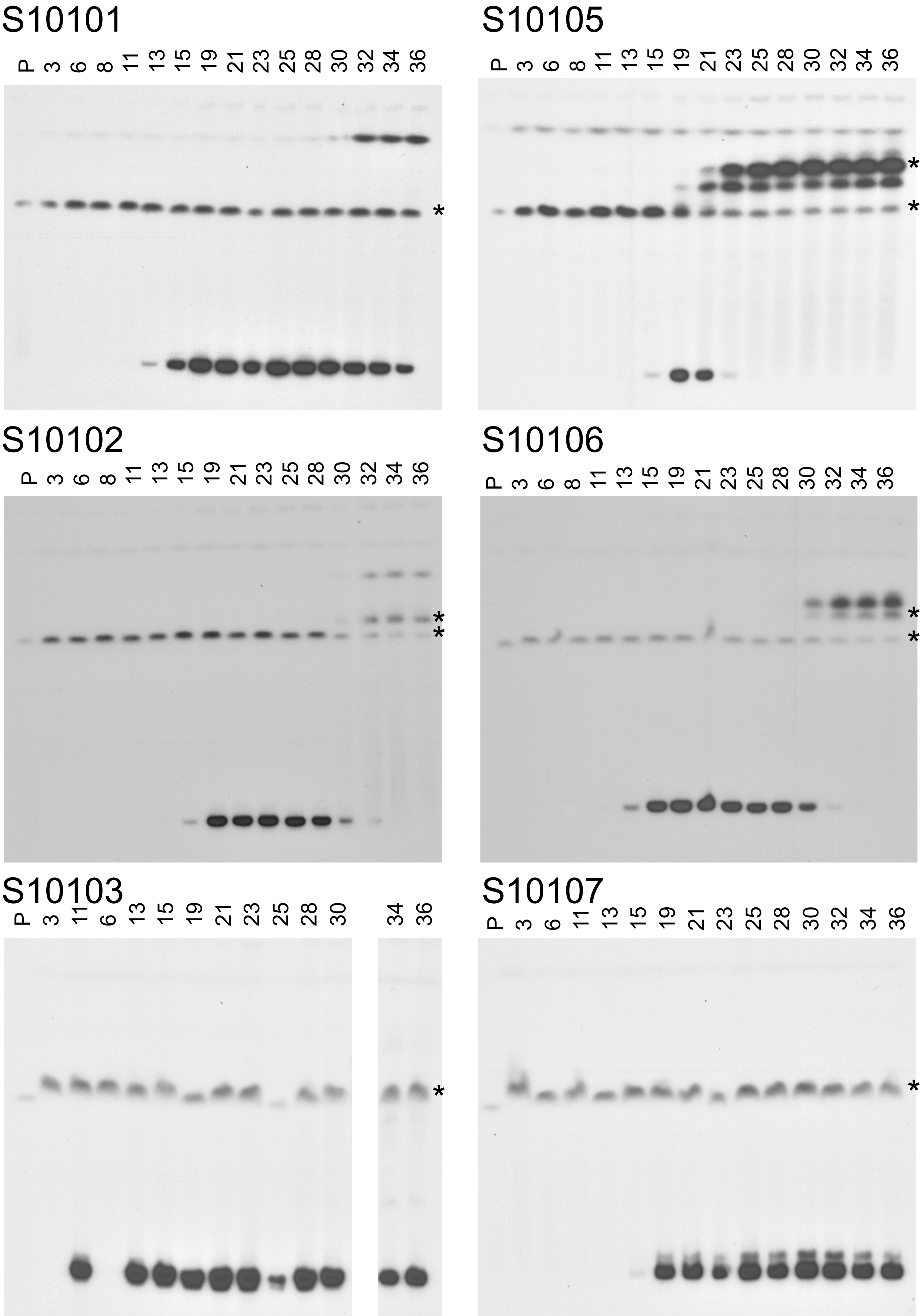
S10101
P 3 6 8 11 13 15 19 21 23 25 28 30 32 34 36
*
S10105
P 3 6 8 11 13 15 19 21 23 25 28 30 32 34 36
*
*
S10102
P 3 6 8 11 13 15 19 21 23 25 28 30 32 34 36
*
*
S10106
P 3 6 8 11 13 15 19 21 23 25 28 30 32 34 36
*
*
S10103
P 3 11 6 13 15 19 21 23 25 28 30 34 36
*
S10107
P 3 6 11 13 15 19 21 23 25 28 30 32 34 36
*

Figure S2. CHEF gel electrophoresis of samples taken at 2-3 day intervals from six chemostat runs containing the heterozygous diploid *ARS228/ars228Δ*. Each CHEF gel was hybridized in turn with *SUL1* (shown) and *CEN2* (*). Each culture acquired extrachromosomal amplicons at some point during their continuous growth. By the end of the experiment, four of the cultures had partially or completely replaced the extrachromosomal amplicons by a chromosomally integrated amplicon that resided either on chromosome II or some other undetermined chromosome. For two cultures, only extrachromosomal amplicons were detected through the end of the experiment.

Figure S3

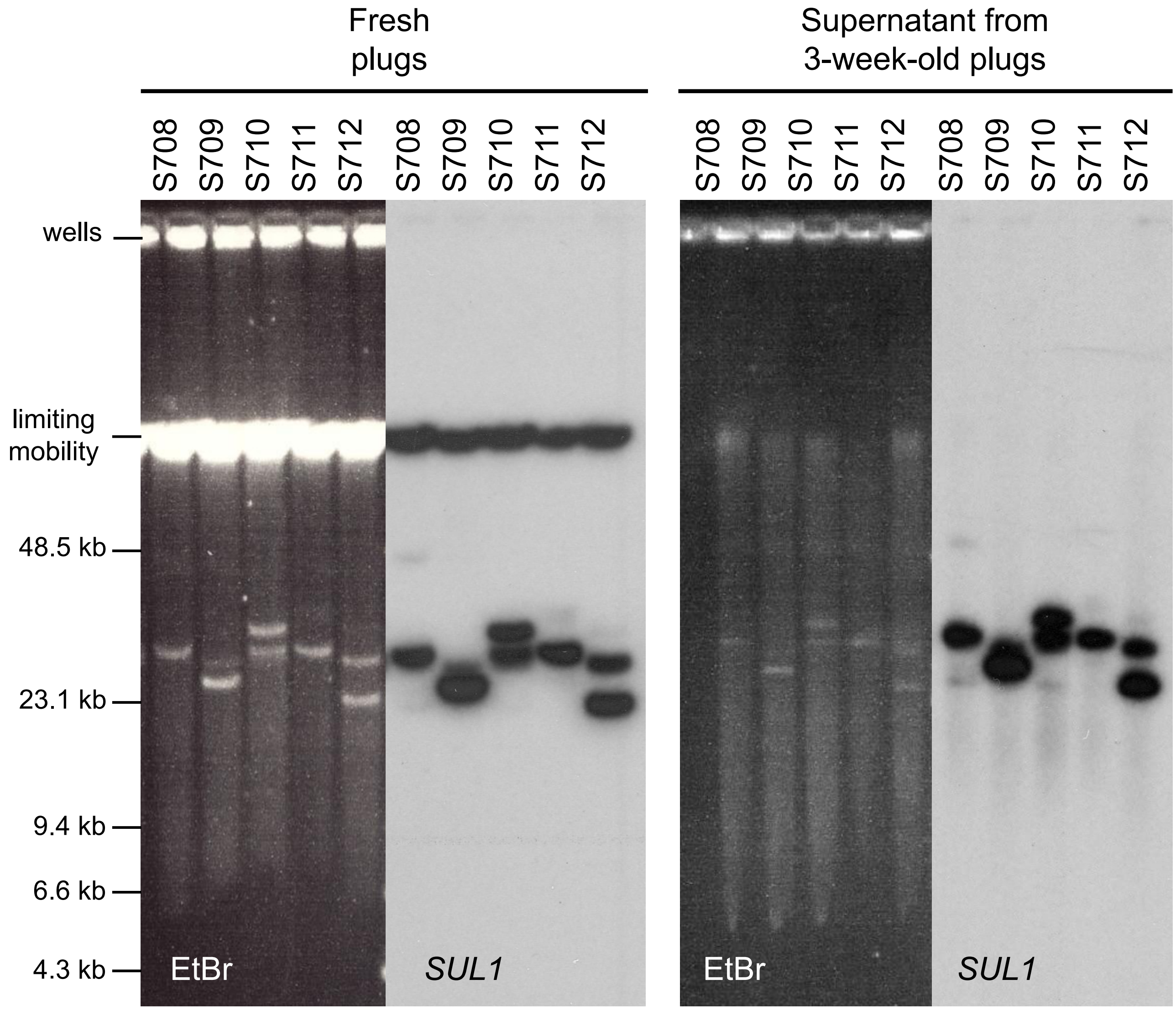


Figure S3. Purification of extrachromosomal amplicons by passive diffusion from agarose plugs. Plugs were prepared from five evolutions on days where extrachromosomal intermediates appeared at their greatest levels. Samples of the freshly made plugs were examined using modified CHEF conditions where all chromosomes run at limiting mobility, but the different versions of extrachromosomal molecules are well separated. After three weeks of passive diffusion, the plug supernatants were collected, concentrated, and run under the same electrophoretic conditions. Both Southern blots were probed with *SUL1*.

Figure S4

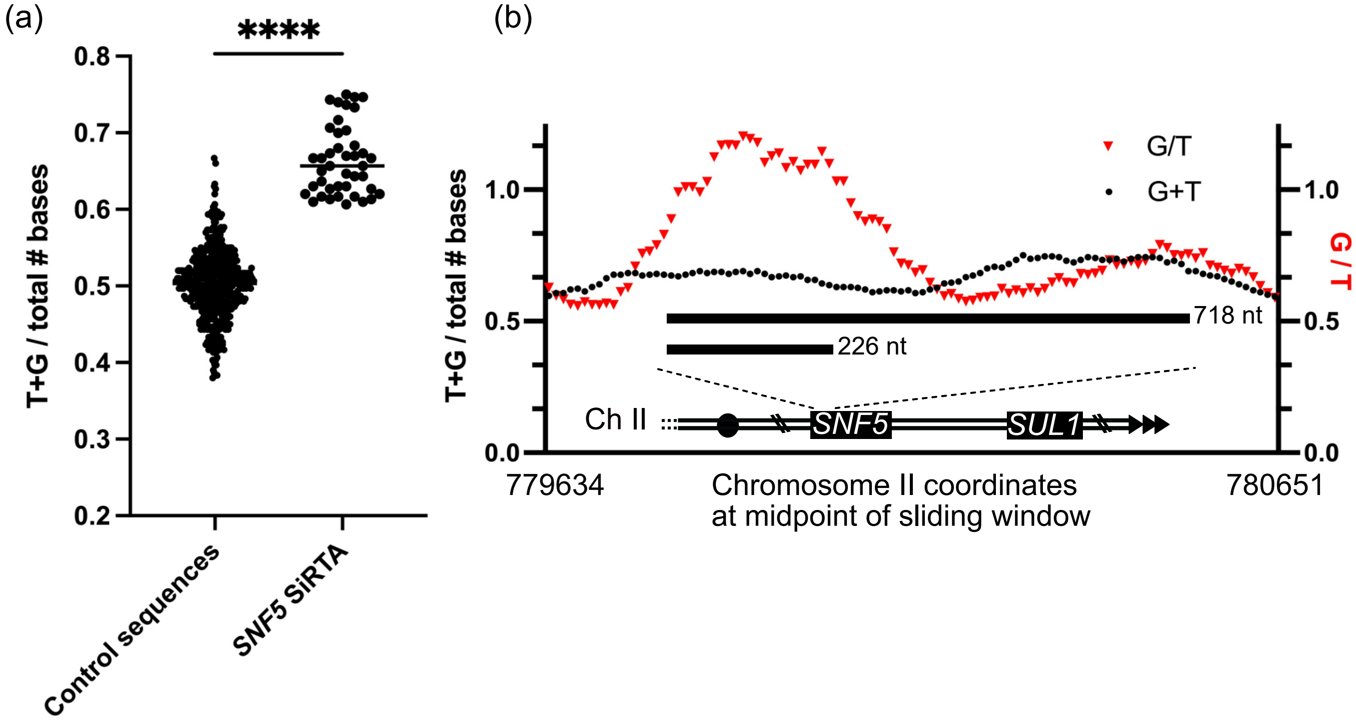


Figure S4. Sequence composition of *SNF5* region. (a) The fraction of thymine and guanine nucleotides (TG-richness) within 493 randomly selected 300 nt sequences is compared to TG-richness of 42 sequences (300 nt each) generated as 10 nt sliding windows across the 718 nt *SNF5* SiRTA. For the *SNF5* SiRTA, only the strand on which telomerase acts to generate a *de novo* telomere is analyzed [the 3' to 5' (bottom) strand of Chromosome II], ***$p < 0.01$ by Mann-Whitney test. (b) The fraction of T and G nucleotides (T+G/300) and ratio of G to T nucleotides (G/T) measured across the *SNF5* SiRTA. 300 nt sequences were generated as 10 nt sliding windows across a region beginning 300 nt 3' to the 718 nt *SNF5* SiRTA and extending 300 nt 5' to the SiRTA. As in (a), only the strand on which telomerase acts to generate a *de novo* telomere is analyzed [the 3' to 5' (bottom) strand of Chromosome II]. The ratio of thymine or guanine among total nucleotides (black) and the ratio of guanine to thymine (red) are shown. Chromosome II coordinates correspond to the midpoint of the read for each data point.

Figure S5

3’**GA|GTGTGGTGTTGTTAACGTCTACTAAGTCGTTGC||TGTGGTTGTCAATGACTCCTCAGCAGA|TG**
**T|TGTTGTT|GTTGTTGTTGTTGTTGTTTGAAGTGGC||GGTGGCGTT|TGCGTAGTTA||GA||||G||**
**|||||GTGG|||GGGAGGAGGCGTTGTTAGA|||GT|||||||||||||||||TGGGTAACGATTAGTTA**
**GTC||GC|||TGAA|GA||||T**G||GGGA||GGAGGAGGAGGTCGTGGTGTGTTGAATGTAGGGGTTTAA
CCAGTTCACGGGAATCGAGGTCGCGGATAATTAAACGGAGGTGTTTAACGAGTCAATGGAAACCGATGTG
TCGTTGTTCAAAACTTGTTCAACTCCGTCGTCCGGTATCGTTTTTTATTAGGTGTCCAACACTTACGTTA
ATGACAACGTGTTGTTGTTCACGTTGCGGTTTAACTCGTCGTTTTCCCTGTCGTTTGCCGTGTTTGAGTC
GATCTTGTCGTCTCCGTTAACGACCAAGTCGTCGTTGTCGTCGTCGTTGAATCTTTGGTTTATGTCGCTG
TTGTCGTTGTTGTCAAATCCGTAGTACACGTTTATGTTGTCGTCGTTGTTTTCGTTGTTGTTGTCGTCGT
CGTCGTAGTCGTTGT|TGTTGTTGTTGTTGTCGTTGTCGTCGTTGTCGTTGTCGTTGTCGTCGTCGTTGT
TGTTGTCGTTGTTGTTGTTGTTGTCGTCGTCGTCGTCGTCGTCGTCGTTCCTGTTTATGGCGTTAGAGTC
GTTCAAGGAG-5’

Figure S5. Comparison of sites of *de novo* telomere addition in the HO experiments with the sulfate limited chemostat experiments. The 175 bp sequence of the TG-rich strand of the *SNF5* SiRTA (3’-5’) is in bold type. Dark blue highlighted lines indicate the position of each telomere addition recovered from chromosome VII. Cyan highlighted lines indicate the positions of each telomere addition recovered from the linear fragments generated in sulfate-limited chemostats.

Figure S6

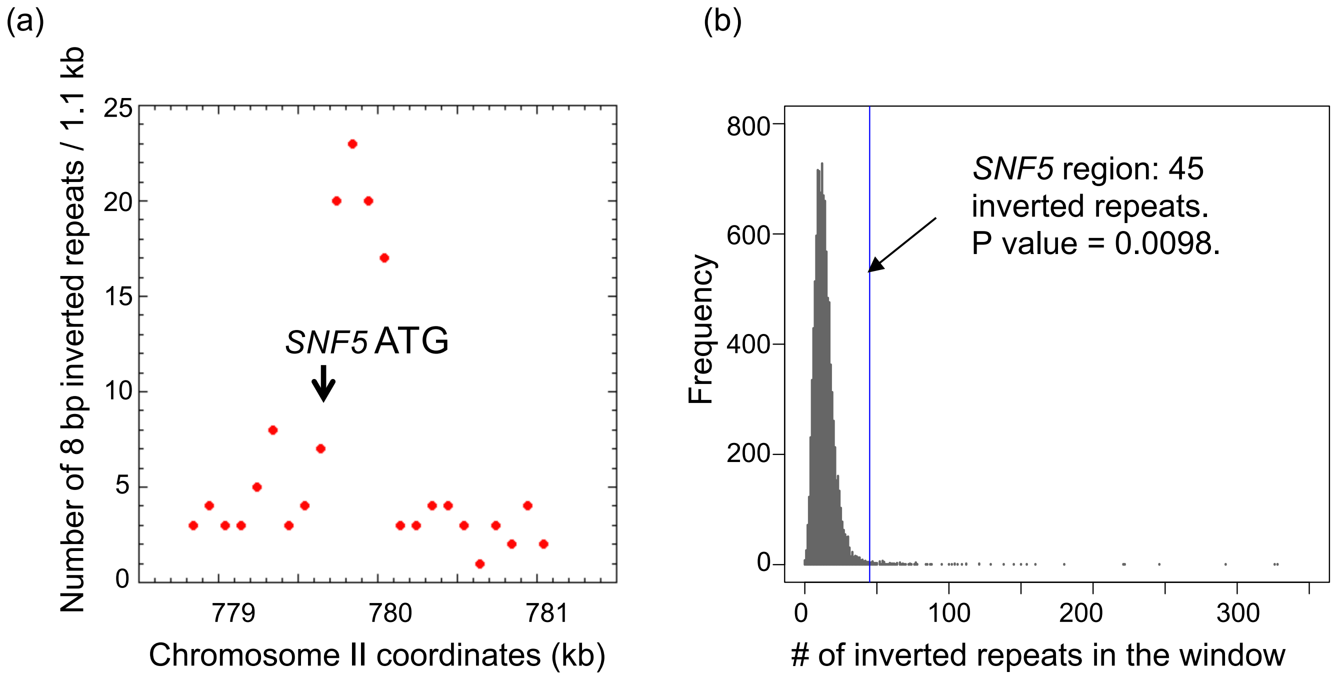


Figure S6. The proposed role of short, interrupted, inverted repeats in the formation of linear *SUL1* amplicons. (a) Overlapping (100 bp step size) 1.18 kb regions containing the majority of the amplicon junctions were scanned for potential interrupted inverted repeats of a minimum of 8 bp separated by ~170 bp or more of intervening sequence. The number of repeat pairs is plotted at the coordinate of the fragment midpoint. (b) As a control we examined the number of 8-mer inverted repeats per 1.18 kb window in non-overlapping windows across the genome (requiring a loop of ≥175 bp). N = 10,309 windows sampled.
